# Ethylene receptors are functionally conserved in calcium permeability across the green lineage

**DOI:** 10.1101/2025.10.20.683334

**Authors:** Dongsheng Yu, Chuanli Ju, Changxin Feng, Yu Wang, Yujia Sun, Lei Gao, Zebin Liu, Chunyan Li, Xuan He, Haimei Su, Mengchen Hu, Yidong Wang, Jie Meng, Shen Tian, Liangyu Liu, Congcong Hou, Dongdong Kong, Legong Li

**Affiliations:** College of Life Sciences, Capital Normal University, and Beijing Key Laboratory of Plant Gene Resources and Biotechnology for Carbon Reduction and Environmental Improvement, Beijing 100048, China

**Keywords:** ethylene receptor, Ca^2+^ permeability, ethylene perception, the plant lineage, evolution

## Abstract

The gaseous hormone ethylene plays a key role in regulating plant growth and stress responses. Although Ca²⁺ has long been implicated in ethylene signaling, the identity of molecules controlling Ca²⁺ permeability involved has remained elusive. Here we show that *Arabidopsis* ethylene receptors ETR1 and ERS1 are Ca^2+^ permeable, and that ETR1 mediates ethylene-induced cytosolic Ca^2+^ increase and regulates hypocotyl elongation. Homologs of ETR1 from eight land plant and algal species also exhibit Ca²⁺ permeability, suggesting an evolutionarily conserved function. We further demonstrate ethylene enhances the Ca^2+^ permeability of ETR1 and its homologue from the charophyte *Klebsormidium flaccidum*. These findings uncover a previously unrecognized but conserved role of ethylene receptors in Ca²⁺ permeability regulation in the green lineage, with broad implications for Ca²⁺ signaling in plant development and environmental adaptation.

## INTRODUCTION

The phytohormone ethylene regulates plant growth and development in response to internal and external cues (Yang and Hoffman, 1984). Plants sense and process the ethylene signal through a pathway that has been extensively studied in the angiosperm *Arabidopsis thaliana*. Ethylene is perceived by five endoplasmic reticulum (ER)-localized ethylene receptors classified into two subfamilies according to sequence features (Wang et al., 2006): subfamily I members, ETHYLENE RESPONSE1 (ETR1) (Chang et al, 1993) and ETHYLENE RESPONSE SENSOR1 (ERS1) (Hua et al., 1995), and subfamily II members, ETR2, ERS2, and ETHYLENE INSENSITIVE4 (EIN4) (Hua et al., 1998; Sakai et al., 1998). In the absence of ethylene, the receptors signal to CONSTITUTIVE TRIPLE RESPONSE 1 (CTR1) (Kieber et al., 1993), a cytosolic serine/threonine protein kinase, which phosphorylates ER-localized ETHYLENE-INSENSITIVE2 (EIN2) (Alonso et al., 1999) to prevent downstream signaling (Ju et al., 2012; Qiao et al., 2012). Ethylene binding inactivates the receptors and CTR1, leading to EIN2 C-terminal cleavage (Ju et al., 2012; Qiao et al., 2012; Wen et al., 2012) and signal transduction in the P-bodies (Li et al., 2015; Merchante et al., 2015) and eventually in the nucleus through EIN3 (Chao et al., 1997; Guo and Ecker, 2003).

Interestingly, the ethylene-signaling pathway combines proteins originating from both prokaryotes and eukaryotes. Ethylene receptors are thought to be derived from the cyanobacterium that underwent endosymbiosis over one billion years ago (Mount and Chang, 2002; Sanderson et al., 2004; Wang et al., 2006), whereas the rest of the key components in the pathway have evolved during the development of the charophyte lineage (Ju et al., 2015). Recently, a functional ethylene receptor, called Synechocystis Ethylene Response1 (SynEtr1), has been demonstrated in the model cyanobacterium (Lacey and Binder, 2016). In addition to plants and cyanobacteria, ETR1 homologs are found in bacteria, as the two-component regulators (Chang et al., 1993), and multiple other microorganisms, as well as a few non-plant eukaryotes (Carlew et al., 2019; Papon and Binder, 2019). Despite the widespread distribution in phylogenetically diverse organisms, the roles of ETR1 and its homologs other than ethylene sensing are largely unclear.

Calcium (Ca²⁺) is a multifunctional second messenger in living organisms (Hetherington and Brownlee, 2004; Kudla et al., 2010), that modulates plant physiological processes through transient fluctuations in the free cytosolic concentration of Ca²⁺ ([Ca^2+^]_cyt_), thereby encoding specific cellular Ca²⁺ signatures (Dodd et al., 2010; Luan and Wang, 2021). The spatiotemporal elevation of [Ca^2+^]_cyt_ is precisely controlled by Ca^2+^-permeable channels and sensed by Ca^2+^ binding proteins, which transmit the signal of the Ca^2+^ transients to initiate downstream biological events (Ward et al., 2009; Demidchik et al., 2018; Tian et al., 2020). Identifying the Ca^2+^ channels that respond to specific stimuli and elucidating their gating mechanisms are central to understanding plant calcium signaling. Over 30 years ago, using the accumulation of the pathogenesis-related protein chitinase as an ethylene response, Raz and Fluhr reported a requirement for Ca^2+^ in ethylene-dependent responses (Raz and Fluhr, 1992). By patch-clamp technique, Zhao et al., (Zhao et al., 2007) examined the effects of ethephon (Warner and Leopold, 1969), an ethylene-releasing compound, and 1-aminocyclopropane-1-carboxylic acid (ACC) (Kende, 1993), an ethylene precursor, on ionic currents in tobacco protoplasts, and the results indicated that ethylene activated a Ca^2+^-permeable channel and thus elicited an elevation of [Ca^2+^]_cyt_. However, the molecules that control the Ca²⁺ permeability associated with ethylene have remained unknown.

Given that calcium channels are membrane proteins, we hypothesized that one or more transmembrane proteins in the *Arabidopsis* ethylene-signaling pathway may possess Ca²⁺ permeability, and we tested this possibility for all five ethylene receptors. Here, we report that the ethylene receptors ETR1 and ERS1, along with their homologs in the green lineage, modulate Ca²⁺ permeability. Pharmacological analyses of the genetic mutants show that ETR1 is required for ethylene-induced increases in [Ca^2+^]_cyt_ and for the regulation of hypocotyl growth in Arabidopsis. We demonstrate the Ca^2+^ conductance of ETR1 and ERS1 using voltage-clamp recordings in *Xenopus laevis* oocytes and reaffirm these results in two additional heterologous expression systems, wherein expression of either protein led to extracellular Ca^2+^-induced [Ca^2+^]_cyt_ increase in human embryonic kidney HEK293T cells and complemented the growth defect of the yeast mutant *cch1 mid1* deficient in Ca^2+^ influx (Fischer et al., 1997). We find also that ethylene modulates the Ca^2+^ permeability of ETR1 and its homolog in algal sister groups, which is strengthened by results from a mutation that disrupts ethylene binding in each protein. Taken together, our findings reveal that ETR1’s Ca²⁺ permeability is broadly distributed across algae and terrestrial plants and suggest that this function, and its regulation by ethylene, originated in a common aquatic ancestor prior to the colonization of land by plants.

## RESULTS

### Ethylene elicits a [Ca^2+^]_cyt_ increase in *Arabidopsis* that requires the involvement of ethylene receptor ETR1

To investigate whether ethylene triggers an increase in [Ca^2+^]_cyt_ in *Arabidopsis*, we used transgenic plants stably expressing aequorin (AQ) (Knight et al., 1991), a bioluminescent reporter for cytoplasmic calcium. Seedlings were treated with ethephon, and [Ca^2+^]_cyt_ changes were monitored. Ethephon application led to a significant elevation in [Ca^2+^]_cyt_ (Figure 1A-1C), but this effect was suppressed by an internal Ca^2+^ channel blocker Ruthenium Red (Knight et al., 1992), to around 70% of the control at 10 µM (Figure 1D and 1E). These data suggest that ethylene-induced [Ca²⁺]_cyt_ increases are mediated at least in part by internal Ca²⁺-permeable channels.

**Figure 1.**
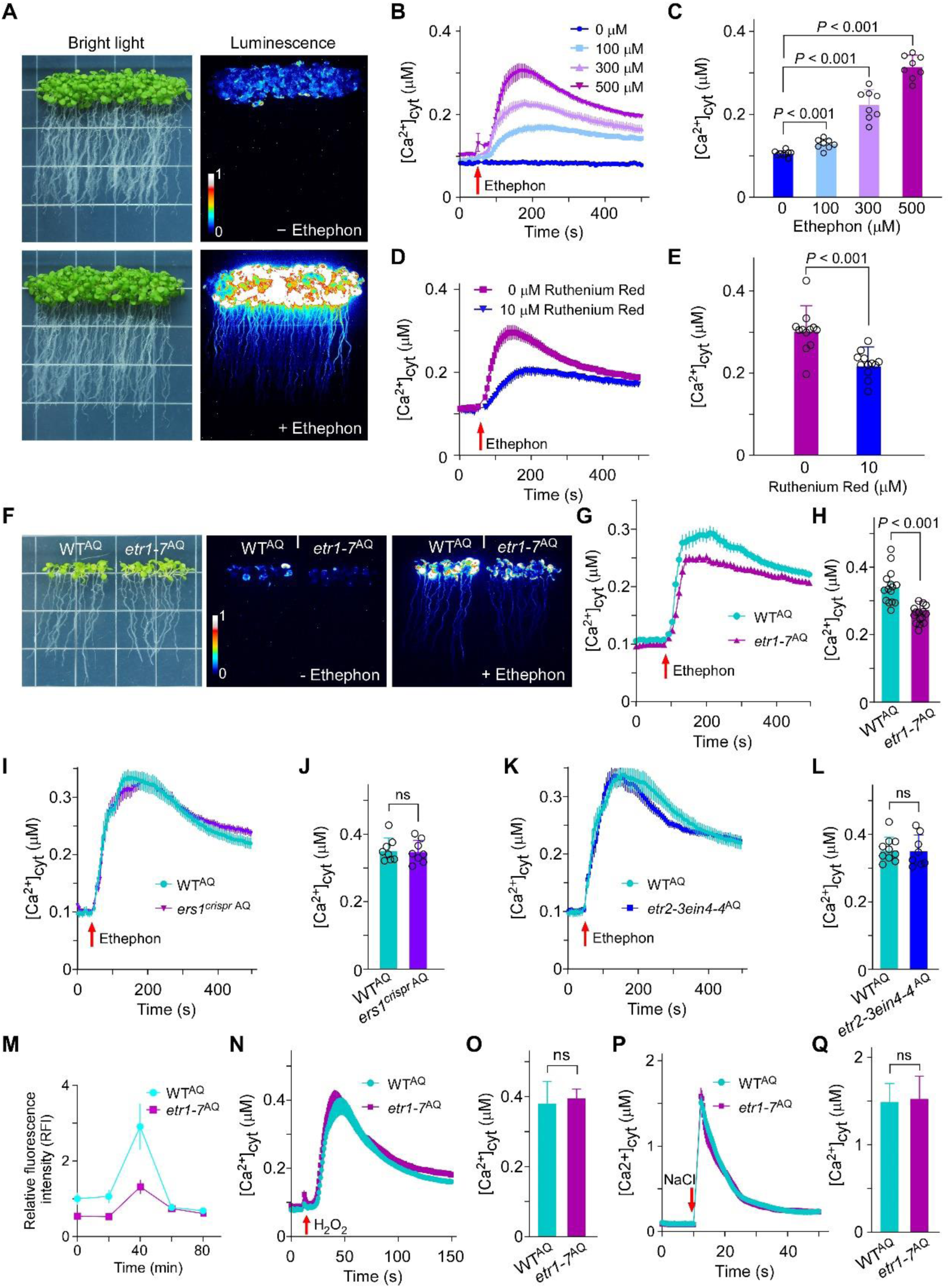
ETR1 is required for ethephon-induced [Ca^2+^]_cyt_ increase in *Arabidopsis*. (**A**) Impact of ethephon on [Ca^2+^]_cyt_ in seven-day-old wild-type (WT) plants stably expressing Ca^2+^-sensitive photoprotein aequorin (WT^AQ^). Representative luminescence images, scaled using a pseudocolour bar, under H_2_O (-ethephon) and ethephon (500 μM, dissolved in water) treatments at the same final pH value (right) and bright-light images before the treatments (left) are shown. (**B**) Time-course of [Ca^2+^]_cyt_ changes in WT^AQ^ seedlings treated with ethephon. Data were obtained using a Varioskan Flash Spectral Scanning Multimode Reader. The arrow indicates the time point of ethephon addition. (**C**) Quantification of [Ca^2+^]_cyt_ in seedlings administered different concentrations of ethephon, based on data in (**B**) when [Ca^2+^]_cyt_ peaked at 500 µM ethephon. The data shown are mean ± s.e., *n* = 8 seedlings. (**D and E**) Influence of Ruthenium Red on ethephon-induced [Ca^2+^]_cyt_ increase. Time-course of [Ca^2+^]_cyt_ changes following treatment with 500 µM ethephon are presented in (**D**), and the data (mean ± s.e., *n* = 8 seedlings) at peak [Ca^2+^]_cyt_ under 0 µM Ruthenium Red are presented in (**E**). In (**D**), the arrow indicates the time point of chemical application. (**F**) Visualization of [Ca^2+^]_cyt_ in WT^AQ^ and the *etr1-7*^AQ^ mutant with (right) and without (middle) ethephon (500 μM) treatments. The bright-light image (left) is also shown. (**G**, **I and K**) Quantification of [Ca^2+^]_cyt_ changes in *etr1-7*^AQ^ (**G**), *ers1*^crisprAQ^ (**I**), and *etr2-3ein4-4*^AQ^ (**K**) mutants over time after ethephon (500 μM) treatment. Data were obtained using a Varioskan Flash Spectral Scanning Multimode Reader. The arrow denotes the time point of ethephon addition. (**H**, **J and L**) Comparison of [Ca^2+^]_cyt_ differences between WT^AQ^ and *etr1-7*^AQ^ (**H**), *ers1*^crisprAQ^ (**J**), and *etr2-3ein4-4*^AQ^ (**L**) seedlings under ethephon treatment, based on the data in (**G**, **I and K**) when WT^AQ^ was at peak [Ca^2+^]_cyt_. (**M**) Quantification of [Ca^2+^]_cyt_ differences between WT^AQ^ and the *etr1-7*^AQ^ mutant over time after ACC (20 μM) treatment. Luminescence data were obtained by aequorin imaging analysis. The arrow indicates the time of ACC addition. (**N and P**) [Ca^2+^]_cyt_ changes in WT^AQ^ and *etr1-7*^AQ^ seedlings in response to H_2_O_2_ (4 mM) (**N**) and NaCl (300 mM) (**P**), based on luminescence data obtained using a Varioskan Flash Spectral Scanning Multimode Reader. (**O and Q**) Comparison of [Ca^2+^]_cyt_ differences between WT^AQ^ and the *etr1-7*^AQ^ mutant under H_2_O_2_ (**O**) and NaCl (**Q**) treatments, based on the data at peak [Ca^2+^]_cyt_ in (**N**) and (**P**), respectively.

We then examined whether any of the known transmembrane proteins in the ethylene-signaling pathway (for example ethylene receptors) is required for increasing ethylene-induced [Ca^2+^]_cyt_ in *Arabidopsis*, and we generated the ethylene receptor single or double knockout mutants expressing aequorin, including *etr1-7*^AQ^, *ers1^crispr^*^AQ^, *etr2-3ein4-4*^AQ^, *etr2-3*^AQ^, and *ein4-4*^AQ^. Compared with the extent of ethephon-increased [Ca^2+^]_cyt_ in WT^AQ^ seedlings, the responses were reduced in *etr1-7*^AQ^ mutants (Figure 1F-1H) but normal in *ers1^crispr^*^AQ^, *etr2-3ein4-1*^AQ^, *etr2-3* ^AQ^, and *ein4-4*^AQ^ mutants (Figure 1I-1L and Supplemental Figures 1 and 2). Besides ethephon, the response of ACC-induced [Ca^2+^]_cyt_ was also reduced in *etr1-7*^AQ^ mutants (Figure 1M and Supplemental Figure 3). By contrast, hydrogen peroxide (H_2_O_2_) (McAinsh et al., 1996) and salt (Knight et al., 1997) elicited a similar [Ca^2+^]_cyt_ fluctuation pattern between WT^AQ^ and *etr1-7*^AQ^ (Figure 1N-1Q). These data suggest that ethylene-stimulated [Ca^2+^]_cyt_ increase in *Arabidopsis* at least partially involves a contribution from ETR1.

### ETR1 and ERS1 are Ca^2+^ permeable

We next tested whether ETR1 or the other ethylene receptors exhibit Ca^2+^ permeability, and we conducted electrophysiological recordings of all five *Arabidopsis* ethylene receptors using *Xenopus laevis* oocytes. GFP fusions confirmed plasma membrane localization of each protein (Supplemental Figure 4), suggesting the feasibility of electrophysiological study in this system. Two-electrode voltage clamp (TEVC) analysis revealed inward Ca²⁺ currents in ETR1-and ERS1-expressing oocytes (Figure 2A and 2B), which were abolished by La^3+^ (Figure 2C and 2D), an external Ca^2+^ channel blocker (Knight et al., 1996). In contrast, we detected no Ca^2+^ currents for ETR2, ERS2, or EIN4 (Figure 2A and 2B). These data show that subfamily I ethylene receptors are Ca^2+^ permeable in the *X*. *laevis* oocyte expression system, but subfamily II ethylene receptors are Ca^2+^ impermeable.

**Figure 2.**
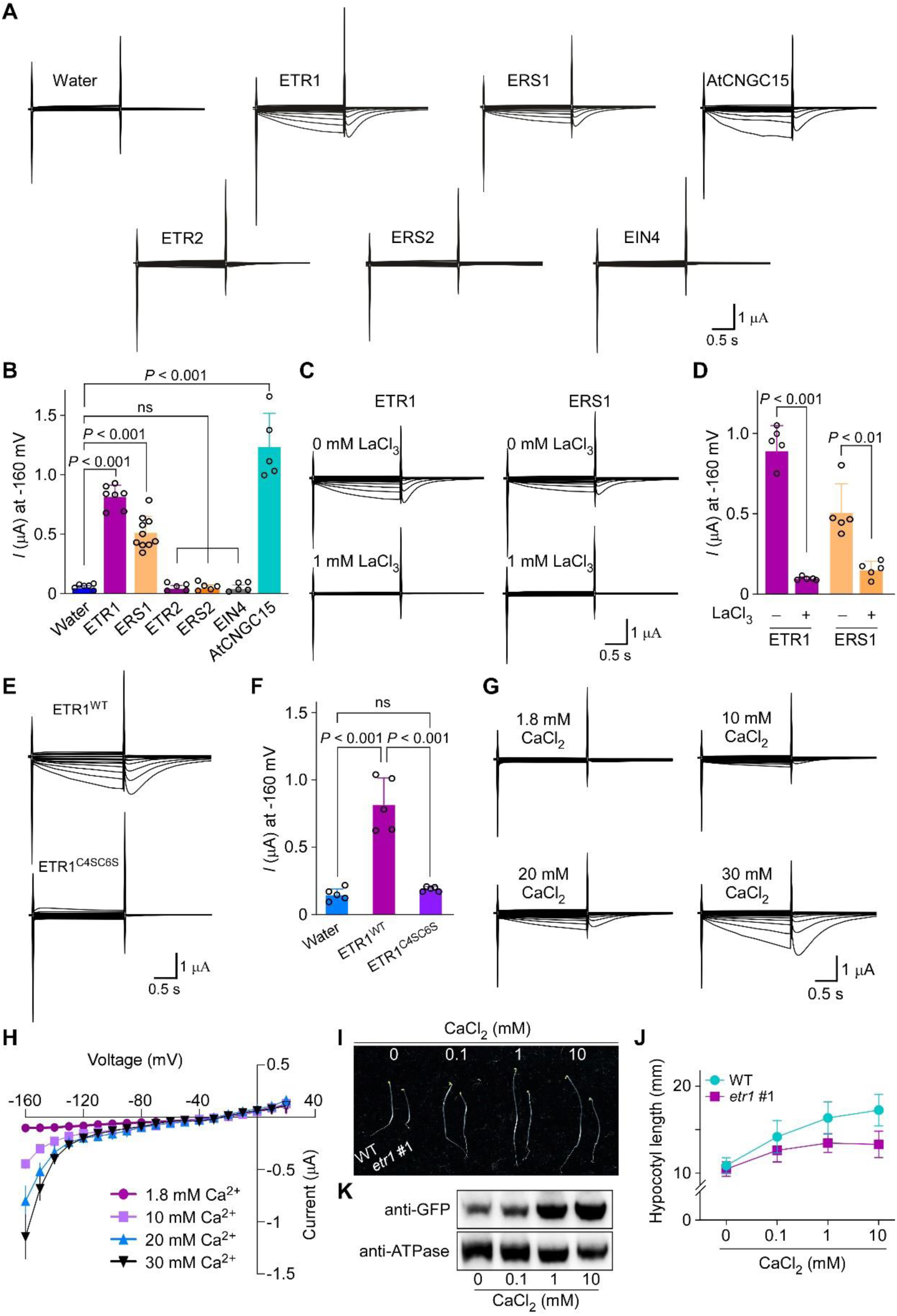
ETR1 and ERS1 are Ca^2+^-permeable in *X. laevis* oocytes, and ETR1 Ca^2+^ permeability governs hypocotyl growth in *Arabidopsis*. (**A**) Representative TEVC recording of Ca^2+^ currents in oocytes expressing ETR1, ERS1, ETR2, ERS2, or EIN4 and perfused with 20 mM Ca^2+^. AtCNGC15 was employed as a positive control. (**B**) Quantification of steady-state ETR1, ERS1, ETR2, ERS2, or EIN4-mediated Ca^2+^ currents at -160 mV based on recordings as in (**A**) (mean ± s.d., *n* ≥ 5 oocytes). (**C and D**) The influence of La^3+^ on ETR1/ERS1-mediated Ca^2+^ currents in oocytes perfused with 20 mM Ca^2+^. Representative current traces without (top panel) and with (bottom panel) LaCl_3_ (1 mM) are presented in (**C)**, and quantified currents are illustrated in (**D)** (mean ± s.d., *n* = 5 oocytes). (**E and F**) Ca^2+^ currents conferred by the WT (ETR1^WT^) and mutant (ETR1^C4SC6S^) variants of ETR1 in oocytes perfused with 20 mM Ca^2+^. ETR1^C4SC6S^ has substituted cysteine at residues 4 and 6 with serine, which disrupts the dimerization between receptor monomers. Representative current traces (**E**) and quantified currents at -160 mV (**F**) (mean ± s.d., *n* = 5 oocytes) are presented. (**G** and **H**) ETR1 Ca^2+^ currents are detected when oocytes are perfused with at least 10 mM Ca^2+^. Representative current traces are presented in (**G**), and the average current-voltage (I-V) relationship is shown in (**H**) (mean ± s.d., *n* = 5 oocytes). (**I**) Representative phenotypes of six-day-old etiolated WT and *etr1* #1 seedlings grown with Ca^2+^. For each comparison, WT is on the left and the *etr1* #1 mutant is on the right. (**J**) Hypocotyl length (mean ± s.d., *n* ≥ 20 seedlings) of genotypes in (**I**). (**K**) Western blot of ETR1-GFP protein under an increased amount of medium Ca^2+^. Six-day-old etiolated WT seedlings stably expressing *ETR1pro::ETR1-GFP* were employed for microsomal fraction isolation. Western blotting was performed using an anti-GFP antibody. ETR1-GFP is approximately 110 kDa. ATPase was utilized as a loading control.

We further validated Ca²⁺ permeability in HEK293T cells and in the yeast *cch1 mid1* mutant, using approaches respectively described for the calcium sensor GCaMP6 (Chen et al., 2013) and MtCNGC15a channel (Charpentier et al., 2016). Upon exposure to elevated extracellular Ca^2+^, more HEK293T cells expressing ETR1 or ERS1 displayed [Ca^2+^]_cyt_ increases compared to the empty vector-transfected controls (Supplemental Figure 5A and 5B). In yeast, ETR1 and ERS1 rescued the pheromone-induced growth defect of the *cch1 mid1* mutant, similar to the positive control, MtCNGC15 (Supplemental Figure 5C). Together, these results support the conclusion that the ethylene receptors ETR1 and ERS1 are Ca^2+^ permeable.

Homodimer is known to be the fundamental unit for ETR1 receptor function (Schaller and Bleecker, 1995; Schaller et al., 1995). To test whether dimerization is required for Ca^2+^ permeability, we mutated the two cysteines (Cys4 and Cys6) at the N terminus of ETR1 to serines to disrupt homodimerization (Schaller and Bleecker, 1995; Schaller et al., 1995). The mutated derivative (ETR1^C4SC6S^) was incapable of mediating Ca^2+^ permeability in contrast with the results obtained with the wild-type (WT) version of ETR1 (ETR1^WT^) (Figure 2E and 2F), which suggests homodimerization is necessary for ETR1 Ca^2+^ permeability. Whether heterodimers or higher-order receptor complexes contribute remains to be determined.

The effect of extracellular Ca^2+^ concentration on ETR1 Ca^2+^ permeability was also measured. Increasing extracellular Ca²⁺ concentrations to 10 mM or higher enhanced ETR1-mediated inward Ca²⁺ currents (Figure 2G and 2H). These results support dose-response Ca²⁺ permeability.

### The Ca^2+^ permeability of ETR1 controls hypocotyl growth

To assess the physiological relevance of ETR1 Ca^2+^ permeability, we analyzed hypocotyl elongation in *etr1* #1 mutant, a T-DNA insertion knockout mutant (Supplemental Figure 6), and WT seedlings. As *etr1* loss-of-function mutants had shorter etiolated hypocotyls (Hua and Meyerowitz, 1998), we examined whether the growth defect in the *etr1* mutant was related to Ca^2+^ by dose-response assays (0, 0.1, 1, and 10 mM). The results in WT seedlings paralleled the medium Ca^2+^ amount, with longer hypocotyls being observed under higher level of Ca^2+^ (Figure 2I and 2J), pointing to a positive role of Ca^2+^ in hypocotyl growth. ETR1 protein levels also increased in stable *ETR1pro::ETR1-GFP* transgenic lines grown under higher Ca^2+^ (Figure 2K). In contrast, the *etr1* #1 mutant exhibited a defect in response to increased amounts of Ca^2+^ (Figure 2I and 2J), indicating impaired Ca^2+^ sensing. These results suggest that ETR1 is essential for Ca^2+^ dependent hypocotyl growth in *Arabidopsis*, potentially explaining earlier reports of ETR1 promoting cell elongation independently of ethylene (Hua and Meyerowitz, 1998).

### ETR1 homologs from the plant lineage are Ca^2+^ permeable

To explore the evolutionary origin of ETR1’s Ca²⁺ permeability, we analyzed ETR1 homologs from diverse plant taxa, including one early diverging alga from Charophyta (*Klebsormidium flaccidum*), two clades within the bryophytes respectively belonging to liverworts (*Marchantia polymorpha*) and mosses (*Physcomitrium patens*), one fern (*Selaginella moellendorffii*), and multiple seed plants classified as dicots (*Brassica napus* and *Medicago truncatula*) and monocots (*Sorghum bicolor* and *Zea mays*) (Figure 3A and Supplemental Figure 7). The cRNA derived from the homolog sharing the highest similarity to ETR1 in each species was individually injected into oocytes for patch-clamp analysis. Clear Ca^2+^ currents were detected for all the ETR1 homologs examined (Figure 3B and 3C). These results suggest that the Ca^2+^ permeability of ETR1 is conserved across land plants and algae and likely originated in an aquatic ancestor of land plants.

**Figure 3.**
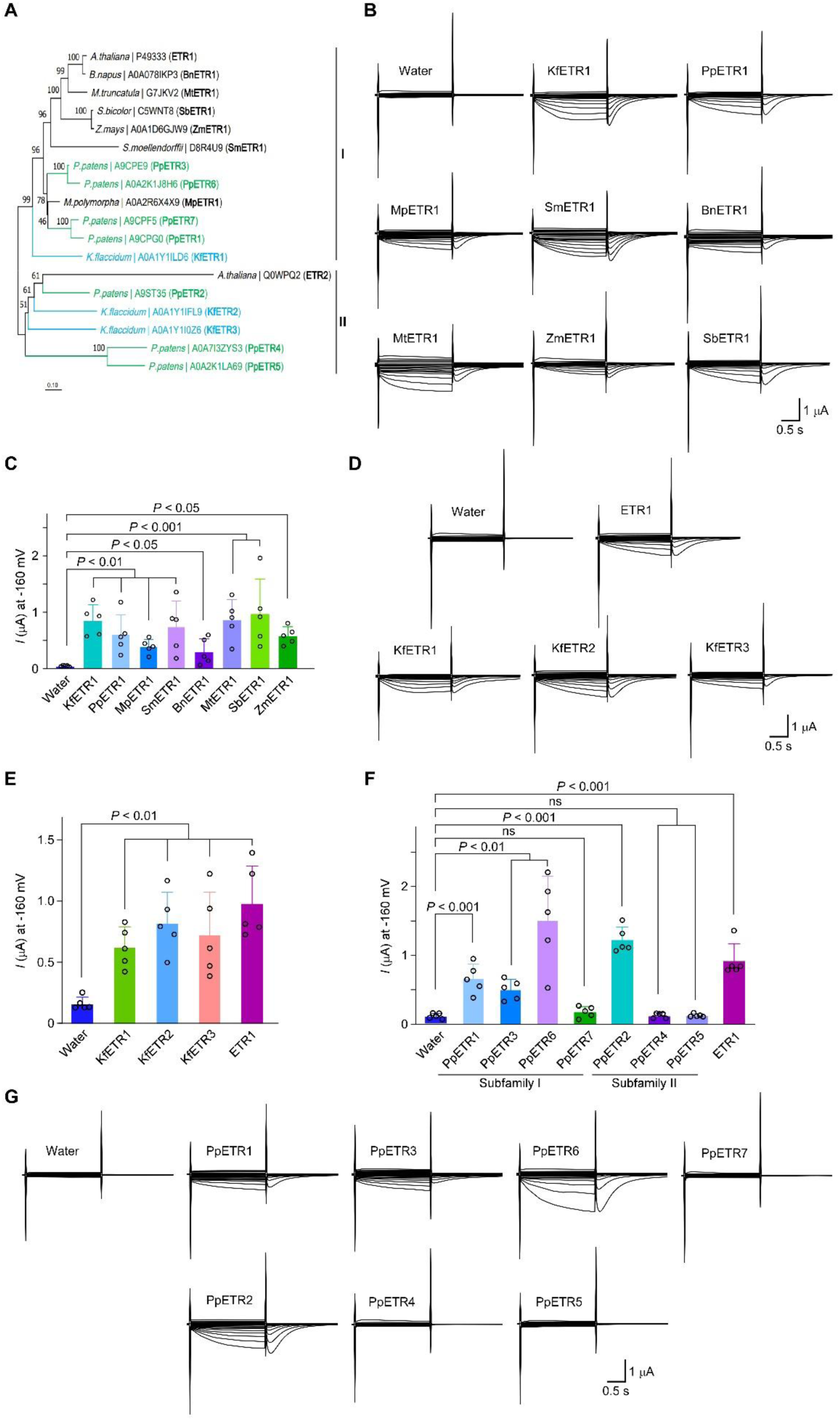
ETR1 homologs from the plant lineage are Ca^2+^-permeable. (**A**) A simplified phylogeny of ETR1 homologs in the plant clade, which was extracted from a wider number of species including plants and non-plants shown in Supplemental Figure 7. The tree presents the relative positions of *Klebsormidium flaccidum* (basal charophyte), *Physcomitrium patens* (moss, bryophyte), *Marchantia polymorpha* (liverwort, bryophyte), *Selaginella moellendorffii* (gymnosperm), *Sorghum bicolor* (monocot angiosperm), *Brassica napus* (eudicot angiosperm), *Medicago truncatula* (eudicot angiosperm), and *Arabidopsis thaliana* (eudicot angiosperm). Protein sequences include all ETR1 homologs in the genomes of *Klebsormidium flaccidum* (blue) and *Physcomitrium patens* (green), as well as the sequence with the highest homology to ETR1 for the rest of the species. Abbreviated names of the species and accession numbers of proteins (https://www.ebi.ac.uk/QuickGO/) are presented. The tree was established using the Neighbor-Joining method, and the numbers adjacent to the branches indicate the bootstrap percentage. (**B and C)** Representative TEVC recording of Ca^2+^ currents in oocytes expressing the homolog most similar to ETR1 from each of the eight species in (**A**) and perfused with 20 mM Ca^2+^. Representative current traces (**B**) and steady-state currents at -160 mV (**C**) (mean ± s.d., *n* = 5 oocytes) are presented. (**D and E)** The Ca^2+^ currents in oocytes expressing each of the ETR1 homologs from *Klebsormidium flaccidum* (KfETRs). ETR1 was employed to compare the currents. Steady-state currents at -160mV (**D**) (mean ± s.d., *n* = 5 oocytes) and representative current traces (**E**) are presented. (**F and G)** Ca^2+^ currents in oocytes expressing each of the ETR1 homologs from *Physcomitrium patens* (PpETRs). ETR1 was employed to compare the currents. Steady-state currents at -160 mV (**F**) (mean ± s.d., *n* = 5 oocytes) and representative current traces (**G**) are presented.

Since only *Arabidopsis* subfamily I ethylene receptors were Ca^2+^ permeable (Figure 2A and 2B), we were curious as to when in plant evolution the Ca^2+^ permeability between ethylene receptor subtypes diverged. Therefore, we examined all ETR1 homologs from *Klebsormidium* (three KfETRs) and *Physcomitrium* (seven PpETRs), which phylogenetically resolve into subfamilies I and II (Figure 3A). All three KfETRs generated positive Ca^2+^ currents (Figure 3D and 3E). In *Physcomitrium*, three of four subfamily I isoforms were permeable, whereas two of three subfamily II isoforms were impermeable (Figure 3F and 3G). These results suggest that subfamily differentiation in Ca²⁺ permeability likely arose after terrestrial colonization.

### Ethylene enhances the Ca^2+^ permeability of ETR1 and KfETR1

Given that ethylene inactivates receptor signaling (Hua and Meyerowitz, 1998), we tested whether it also affects Ca²⁺ permeability. We found that ethephon treatment further increased the Ca^2+^ currents of ETR1^WT^ in oocytes (Figure 4A and 4B). Although a mutation (Cys65Ser) that disrupts ethylene binding in ETR1 (Wang et al., 2006; Schaller and Bleecker, 1995) (ETR1^C65S^) had no obvious effect on the Ca^2+^ current size under zero ethephon conditions, the ability of ethephon to elevate the Ca^2+^ current was abolished (Figure 4A and 4B), indicating a ligand dependent mechanism.

**Figure 4.**
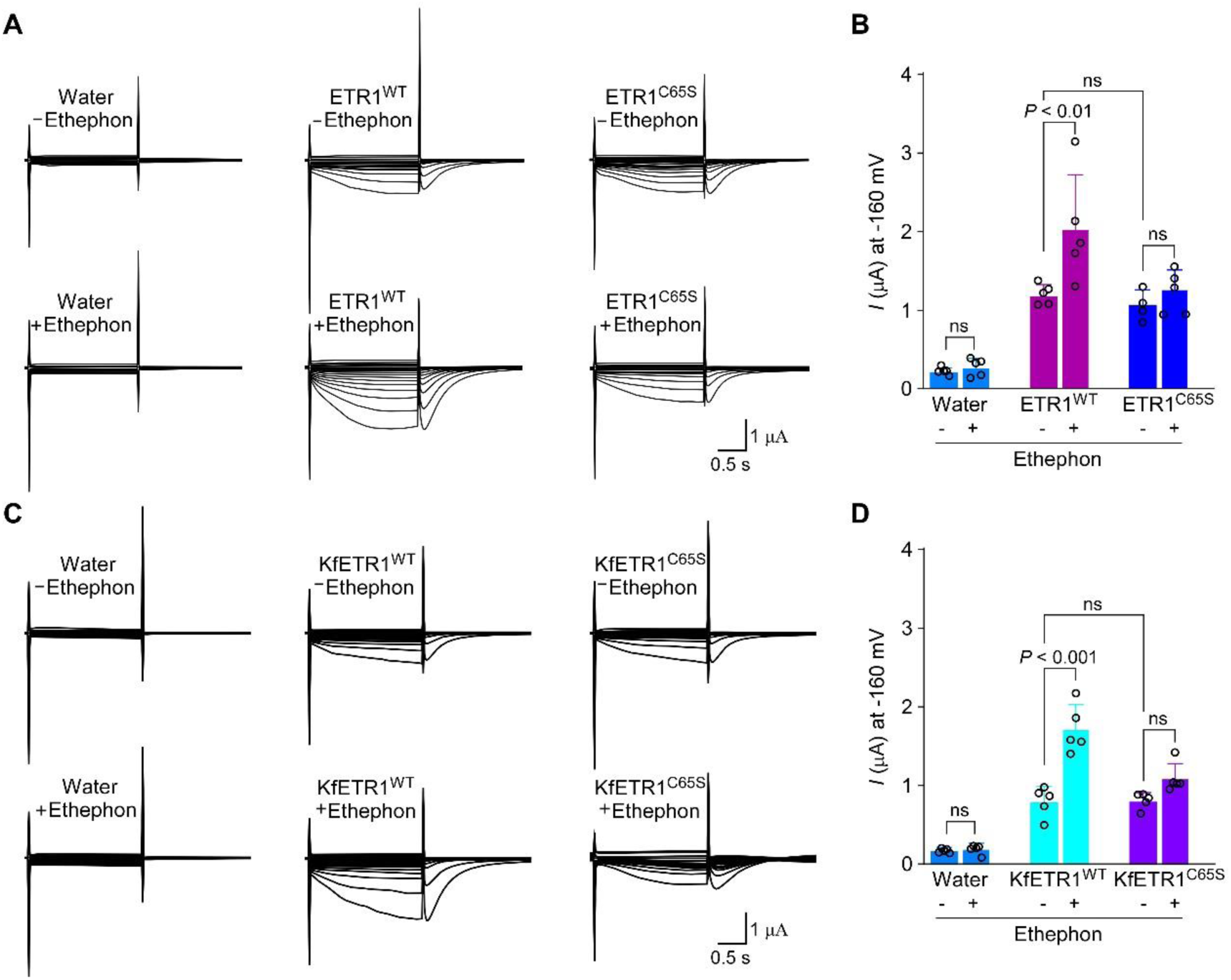
Ethylene increases the Ca^2+^ permeability of ETR1 and its homolog from *Klebsormidium*. **(A and B)** Comparison of Ca^2+^ currents conferred by ETR1^WT^ and ETR1^C65S^ with (bottom panel) and without (top panel) ethephon in oocytes perfused with 20 mM Ca^2+^. ETR1^C65S^ contains a cysteine-to-serine substitution at residue^65^, blocking ethylene binding to the receptor. Representative current traces (**A**) and quantified steady-state currents at -160 mV (**B**) (mean ± s.d., *n* = 5 oocytes) are presented. (**C and D)** Comparison of Ca^2+^ currents conferred by KfETR1^WT^ and KfETR1^C65S^ without (top panel) and with (bottom panel) ethephon (500 μM) in oocytes perfused with 20 mM Ca^2+^. A cysteine-to-serine mutation was introduced in KfETR1 at residue^65^ of ETR1 to create KfETR1^C65S^ for Ca^2+^ current characterization in oocytes. Representative current traces (**C**) and steady-state currents at -160 mV (**D**) (mean ± s.d., *n* = 5 oocytes) are shown.

We further tested whether the ethylene regulation of ETR1 Ca^2+^ permeability is conserved in algae. We found that ethephon enhanced the Ca^2+^ permeability of KfETR1^WT^, and mutation of the conserved cysteine residue identical to Cys^65^ in ETR1 (KfETR1^C65S^) (Supplemental Figure 8) abolished the ethephon effect (Figure 4C and 4D). These data suggest that ethylene-mediated modulation of ETR1 function predates land plant evolution.

## DISCUSSION

Although the presence of ethylene-responsive Ca^2+^-permeable channels has long been hypothesized in plants, none had been identified. The current work reveals ethylene receptors ETR1 and ERS1 are Ca^2+^ permeable and ETR1 modulates hypocotyl growth and ethylene-induced [Ca^2+^]_cyt_ increases in *Arabidopsis*. Furthermore, we show that this Ca^2+^ permeability is conserved in ETR1 homologs across algae and land plants, indicating a dual function, ethylene perception and Ca²⁺ conductance, that likely emerged before the emergence of land plants. Our findings also highlight the prominence of metal ions, on the basis of previously reported copper (Hirayama et al., 1999) and silver (McDaniel et al., 2012), in plant ethylene response.

Intriguingly, both ETR1 and KfETR1 were Ca^2+^ permeable in the absence and presence of ethylene. This contrasts with the conventional ON/OFF receptor model, where ethylene inactivates signaling. Instead, our results suggest ethylene enhances ETR1’s Ca²⁺ permeability, thereby reinforcing downstream Ca²⁺ signals. Thus, ETR1 appears to act as a channel that is modulated, rather than silenced, by ethylene. These findings may also highlight a potential general involvement of these ETRs, as Ca^2+^ channels, in plant developmental events and stress responses during evolution, under which conditions transient [Ca^2+^]_cyt_ fluctuations frequently occur.

Besides being a vital hormone, ethylene is also a volatile organic compound (VOC) (Laothawornkitkul et al., 2009; Loreto and D’Auria, 2022). In animals, VOCs are detected by odorant-binding proteins (OBP) and olfactory receptors (ORs), which trigger Ca²⁺ influx and downstream responses via ion channels (Buck, 1996; Loreto and D’Auria, 2022). To date, no plant Ca²⁺ channels have been implicated in VOC signaling (Loreto and D’Auria, 2022). Our findings imply that plants may bypass the need for an animal-like OBP-ORs molecular network to modulate [Ca^2+^]_cyt_, instead relying on ethylene receptors themselves to mediate VOC-triggered Ca²⁺ signals.

The finding ETR1 and ERS1 facilitate Ca^2+^ permeability emphasizes the advantages of localizing ethylene receptors to the ER membrane. We propose that freely diffusible nature of ethylene enables binding to ETR1 and ERS1, and at the same time it can trigger Ca²⁺ release from internal ER stores thereby leading to [Ca^2+^]_cyt_ elevation. In addition to ER-based Ca^2+^ release via ETR1/ERS1, the ethylene-induced [Ca^2+^]_cyt_ increase we observed in Arabidopsis could also arise from Ca^2+^ influx across the plasma membrane via unknown channels, as La^3+^ repressed part of the [Ca^2+^]_cyt_ increase (Supplemental Figure 9). Future work to discover these channels and identify the downstream decoding components of ETR1-mediated Ca^2+^ signal will draw a more complete picture of ethylene-induced Ca^2+^ signature.

Our data showed that in *Arabidopsis* only subfamily I ethylene receptors are Ca^2+^-permeable, whereas in more ancient lineages, *Klebsormidium* and *Physcomitrium*, the Ca^2+^ permeability occurs in both ethylene receptor subfamilies. The emergence of this functional split after land colonization, evident in *Physcomitrium*, may relate to differences in receptor structure or kinase activity. Our data from oocytes showed that REVERSION-TO-ETHYLENE SENSITIVITY1 (RTE1) (Resnick et al., 2006), a protein works upstream of ETR1 and seems specific to ETR1 (Resnick et al., 2008), had no influence on ETR1 Ca^2+^ permeability (Supplemental Figure 10). Exploration of the relationship between Ca^2+^ permeability and ethylene responses may help understand the most fundamental difference between ethylene receptor subfamilies, from the perspective of the Ca^2+^ permeability function.

Our finding that KfETR1 is Ca^2+^ permeable implies an origin of this function prior to the emergence of terrestrial plants. Investigating the Ca^2+^ permeability of ETR1 homologs in eubacteria, for instance SynEtr1, and fungi as well as non-plant eukaryotes, which lack ethylene-binding activity (Wang et al., 2006), may shed light on the evolutionary pressures shaping this dual functionality and may provide a wide perspective on ETR1 function in evolution.

## METHODS

### Plant materials and growth conditions

The *Arabidopsis thaliana* ecotype Columbia-0 (Col-0) was used as the wild type (WT) in this study. *35S::aequorin* seeds (WT^AQ^) were provided by Prof. Marc Knight (Durham University). The *etr1 #*1 mutant, a T-DNA ETR1 (SALK_136293) insertion line, was obtained from the *Arabidopsis* Biological Resource Center (ABRC) and genotyped using primers listed in Supplemental Table1. The *etr1-7*^AQ^, *etr2-3ein4-4*^AQ^, *etr2-3* ^AQ^, and *ein4-4*^AQ^ mutants were obtained from the F2 progeny of a genetic cross between the *etr1-7 etr2-3 ein4-4* triple ethylene receptor mutant and a WT^AQ^ plant. The homozygosity for three ethylene receptors was validated through DNA sequencing, and the aequorin loci were confirmed by genotyping with primers listed in Supplemental Table1. The *ers1^crispr^* mutant was created from CRISPR/Cas9 approach by using pHEN401 expression vector. The constructed plasmid was transformed into Col-0 WT plants by the Agrobacterium (GV3101)-mediated floral-dip method. T2 homozygous plants were underwent PCR-based genotyping to verify the homozygosity and the absence of CRISPR/Cas9. The *ers1^crispr^*^AQ^ material was selected from the F2 progeny of a genetic cross between *ers1^crispr^*mutant and WT^AQ^ plant. For dark-grown seedling phenotypes, sterilized seeds were planted on MS-based calcium-free medium supplemented with specified concentrations of calcium and 0.8% agar. After 3 days of culturing at 4 ℃, the seeds were grown under light for 5-6 hours and incubated in the dark for 6 days at 22 ℃. Hypocotyl length was determined using ImageJ software (http://rsbweb.nih.gov/ij/) on digital images. Three biological replicates were conducted for both *etr1 #*1 and *etr1-7^AQ^* mutants with similar result patterns, and the data from *etr1 #*1 mutant are presented. To prepare seedlings for [Ca^2+^]_cyt_ measurement, sterilized seeds were grown on half-strength MS medium and exposed to white light in a growth chamber at 22 ℃ with a 16-hour light/8 8-hour dark photoperiod for 7 days.

### Visualization of [Ca^2+^]_cyt_ in *Arabidopsis* by aequorin imaging

The 7-day-old *Arabidopsis* WT^AQ^ and *etr1-7*^AQ^ mutant seedlings were sprayed with 10 μM coelenterazine (Nanolight) and placed in the dark for 8 hours before analysis. Aequorin bioluminescence was recorded by the Tanon 5500 Multi Chemiluminescent Imaging System, and bioluminescence images were examined using ImageJ. A 5-minute recording of luminescence images was performed both before the indicated treatment to show [Ca^2+^]_cyt_ at the resting state and 1 minute after the chemical application to detect [Ca^2+^]_cyt_ changes. The total aequorin amount was evaluated by the treatment of the seedlings using discharging buffer [2 M CaCl_2_ in 20% (v/v) ethanol]. The [Ca^2+^]_cyt_ levels were calibrated according to luminescence values as described previously.

### Two-electrode voltage clamp (TEVC) recording in *Xenopus laevis* oocytes

Full-length CDSs of target genes, obtained via PCR amplification of *Arabidopsis* cDNAs with primers listed in Supplemental Table1 or via chemical synthesis, were cloned into the pGEMHE vector. Using the linearized plasmid as the template, capped RNA (cRNA) was produced according to the mMESSAGE mMACHINE T7 kit (Ambion) instructions. Following the injection of 23 ng of cRNA into each oocyte, the oocytes were incubated at ND96 solution (96 mM NaCl, 2 mM KCl, 1 mM MgCl_2_, 1.8 mM CaCl_2_, and 10 mM HEPES, pH 7.4) containing 50 μg/mL gentamicin at 18 ℃ for two days. The standard solution used for TEVC recording contained 2 mM NaCl, 2 mM KCl, 1 mM MgCl_2_, 1.8 mM CaCl_2_, and 10 mM HEPES (pH 7.4), with an osmolality of 220 mOsmo/kg adjusted using mannitol. Oocytes were voltage-clamped using a TEV 200A amplifier (Dagan) and monitored with a computer equipped with a Digidata 1440A converter (Axon). Electrophysiological recording was performed using pCLAMP 10.7 software (Axon). The voltage tested ranged from +20 to -160 mV (in -10 mV decrements, 2-s duration), with a holding potential of -30 mV. For ion selectivity analysis, the perfusion solution contained 10 mM HEPES (pH 7.4) and 15 mM divalent or 30 mM monovalent cation.

### Subcellular localization of target proteins in *X. laevis* oocytes

The full-length CDS of the *Arabidopsis* gene fused with GFP was constructed into the pGEMHE vector using primers listed in Supplemental Table1. The cRNA was prepared using a mMESSAGE mMACHINE T7 kit (Thermo Fisher). Following an injection of the cRNA into oocytes, the cells were incubated at 18 ℃ for two days. Imaging of GFP was performed using a confocal laser scanning microscope (Leica TCS SP8).

### Western blot analysis

Stable *ETR1pro::gETR1-GFP* transgenes were produced for ETR1 protein detection. Using genomic DNA from *Arabidopsis* WT as the template, a 5 kb fragment (including 2.3 kb upstream of the start codon of *ETR1* and the full-length genomic sequence of *ETR1* lacking the stop codon) was PCR amplified using primers listed in Supplemental Table1 and cloned into the pCAMBIA1300-GFP vector. The obtained *ETR1pro::gETR1-GFP* recombinant plasmid was transformed into *Arabidopsis* WT via the *Agrobacterium*-mediated inflorescence dipping method. Transformants with a single insertion were selected using hygromycin. Homozygous lines in the T3 generation were used for analysis.

Total protein was extracted at 4 °C using extraction buffer [50 mM Tris (pH 8.5), 150 mM NaCl, 10 mM EDTA, 20% glycerol (v/v), and protease inhibitors]. After centrifugation at 8,000 × *g* for 15 minutes at 4 °C, the supernatant was collected and centrifuged at 100,000 × *g* for 30 minutes. The membrane pellet was resuspended in resuspension buffer [10 mM Tris (pH 7.5), 150 mM NaCl, 1 mM EDTA, 10% glycerol (v/v), 0.4% (w/v) sodium deoxycholate, and protease inhibitors]. The concentration of the microsomal fraction was determined using a BCA assay kit. The protein was separated via SDS-PAGE (8%) and transferred to a PVDF membrane via electroblotting. Immunoblotting was performed with an anti-GFP antibody (Abclonal) and then with an anti-α-ATPase antibody (PhytoAB) after stripping the membrane.

### Imaging of extracellular Ca^2+^-induced [Ca^2+^]_cyt_ increase in HEK293T cells

The fluorescent calcium indicator GCaMP6s was utilized to monitor [Ca^2+^]_cyt_ in HEK293T. The full-length CDS of the gene was inserted into the pBudCE4.1-GCaMP6s vector using the primers listed in Supplemental Table1. The recombinant plasmid and empty vector were transfected into HEK293T cells, and the cells were cultured in DMEM medium supplemented with 10% FBS under 5% CO_2_ at 37 ℃ for 18-24 hours. For Ca^2+^-induced [Ca^2+^]_cyt_ increase recording, the cells were cultured in standard buffer [120 mM NaCl, 3 mM KCl, 1 mM MgCl_2_, 1.2 mM NaHCO_3_, 10 mM glucose, 10 mM HEPES (pH 7.4)] with 0.1 mM CaCl_2_ for 30 minutes and then in standard buffer with 2.5 mM CaCl_2_. Ca^2+^ imaging was performed using a Leica SP8 microscope, and fluorescence signals were collected every 5 seconds, starting 60 seconds before the changes of bath Ca^2+^ to 2.5 mM.

### Complementation in growth phenotypes of the yeast *cch1 mid1* mutant

The coding sequence of the target gene was PCR amplified using primers listed in Supplemental Table1 and cloned into the YEP124 vector via homologous recombination. The yeast WT strain (BY4741) and the *mid1 cch1* mutant were used as previously reported. The recombinant plasmid and YEP124 empty vector were transformed into the *mid1 cch1* mutant, and transformants were selected on YPD medium without uracil. Yeast growth was examined as described. The yeast was spread evenly on YPD agar medium, and sterilized cellulose filter discs (in 6 mm diameter) pre-soaked with 10 µg synthetic ɑ factor (Sigma) were placed on the medium. After yeast growth at 28 °C for 48 hours, pictures were taken. The yeast mutant complemented with *MtCNGC15a* was employed as a positive control.

### Recording of [Ca^2+^]_cyt_ bursts in *Arabidopsis* following chemical treatment

WT^AQ^ and *etr1-7*^AQ^, *ers1^crispr^*^AQ^, *etr2-3ein4-4*^AQ^, *etr2-3* ^AQ^, *ein4-4*^AQ^ seedlings were sprayed with coelenterazine (10 μM) and kept in the dark overnight. Kinetic bioluminescence was monitored at intervals of 0.25 seconds or 5 seconds using a Varioskan Flash Spectral Scanning Multimode Reader (Thermo Fisher). After a 30-second recording of the resting [Ca^2+^]_cyt_ level, bioluminescence values were obtained for 50-400 seconds to illustrate ethephon (or other indicated chemical)-induced [Ca^2+^]_cyt_ increase. Luminescence values were used for [Ca^2+^]_cyt_ calibration as described previously.

### Gene expression analysis

Total RNA was isolated from the indicated plants using TRIzol (Invitrogen) and reverse-transcribed using Superscript V (Invitrogen) following the manufacturer’s instructions. cDNAs from WT and *etr1* #1 mutant were used as templates for gene expression analysis using primers listed in Supplemental Table1. *Arabidopsis ACTIN* was employed as an internal reference for normalization. Three biological replicates were conducted, and the results from one replicate are shown.

### Statistical analysis

Statistical analyses were performed using two-way ANOVA or two-sided Student’s *t*-tests. Significant difference between samples was determined at the criterion of *P* ≤ 0.05, with ns (not significant) representing *P* > 0.05.

## AUTHOR CONTRIBUTIONS

D.K., L.G.L., and C.J. conceived the project and designed the research. D.Y., C.J., C.F., Y.W., Y.S., L.G., Z.L., C.L., X.H., H.S., M.H., Y.D.W. and J.M. performed the experiments, D.Y., C.J., C.F., Y.W., Y.S., L.G., Z.L., C.L., X.H., H.S., M.H., Y.D.W., J.M., S.T., L.Y.L., C.H., D.K., and L.G.L. participated in the data analysis. C.J., D.K., and L.G.L. wrote the manuscript with contributions from all authors. All authors contributed to the article discussed the results and have approved the submitted version.

## ACKNOWLEDGMENTS

We thank Dr. Caren Chang (University of Maryland) for providing helpful comments on the manuscript. The authors declare no conflict of interest.

## REFERENCES

1. Alonso, J. M., Hirayama, T., Roman, G., Nourizadeh, S., Ecker, J. R. (1999). EIN2, a bifunctional transducer of ethylene and stress responses in Arabidopsis. Science 284: 2148–2152.

2. Buck, L. B. (1996). Information coding in the vertebrate olfactory system. Annu. Rev. Neurosci. 19: 517–544.

3. Carlew, T. S., Allen, C. J., Binder, B. M. (2019). Ethylene Receptors in Nonplant Species. Small Methods 1900266.

4. Chang, C., Kwok, S. F., Bleecker, A. B., Meyerowitz, E. M. (1993). Arabidopsis ethylene-response gene ETR1: similarity of product to two-component regulators. Science 262: 539–544.

5. Chao, Q., Rothenberg, M., Solano, R., Roman, G., Terzaghi, W., Ecker, J. R. (1997). Activation of the ethylene gas response pathway in Arabidopsis by the nuclear protein ETHYLENE-INSENSITIVE3 and related proteins. Cell 89: 1133–1144.

6. Charpentier, M., Sun, J., Vaz Martins, T., Radhakrishnan, G. V., Findlay, K., Soumpourou, E., Thouin, J., Véry, A. A., Sanders, D., Morris, R. J., et al. (2016). Nuclear-localized cyclic nucleotide-gated channels mediate symbiotic calcium oscillations. Science 352: 1102–1105.

7. Chen, T. W., Wardill, T. J., Sun, Y., Pulver, S. R., Renninger, S. L., Baohan, A., Schreiter, E. R., Kerr, R. A., Orger, M. B., Jayaraman, V., et al. (2013). Ultrasensitive fluorescent proteins for imaging neuronal activity. Nature 499: 295–300.

8. Demidchik, V., Shabala, S., Isayenkov, S., Cuin, T. A., Pottosin, I. (2018). Calcium transport across plant membranes: mechanisms and functions. New Phytol. 220: 49–69.

9. Dodd, A. N., Kudla, J., Sanders, D. (2010). The language of calcium signaling. Annu. Rev. Plant Biol. 61: 593–620.

10. Fischer, M., Schnell, N., Chattaway, J., Davies, P., Dixon, G., Sanders, D. (1997). The *Saccharomyces cerevisiae* CCH1 gene is involved in calcium influx and mating. FEBS Lett. 419: 259–262.

11. Guo, H., and Ecker, J. R. (2003). Plant responses to ethylene gas are mediated by SCF(EBF1/EBF2)-dependent proteolysis of EIN3 transcription factor. Cell 115: 667–677.

12. Hetherington, A. M., and Brownlee, C. (2004). The generation of Ca(2+) signals in plants. Annu. Rev. Plant Biol. 55: 401–427.

13. Hirayama, T., Kieber, J. J., Hirayama, N., Kogan, M., Guzman, P., Nourizadeh, S., Alonso, J. M., Dailey, W. P., Dancis, A., Ecker, J. R. (1999). RESPONSIVE-TO-ANTAGONIST1, a Menkes/Wilson disease-related copper transporter, is required for ethylene signaling in Arabidopsis. Cell 97: 383–393.

14. Hua, J., Chang, C., Sun, Q., Meyerowitz, E. M. (1995). Ethylene insensitivity conferred by Arabidopsis ERS gene. Science 269: 1712–1714.

15. Hua, J., and Meyerowitz, E. M. (1998). Ethylene responses are negatively regulated by a receptor gene family in *Arabidopsis thaliana*. Cell 94: 261–271.

16. Hua, J., Sakai, H., Nourizadeh, S., Chen, Q. G., Bleecker, A. B., Ecker, J. R., Meyerowitz, E. M. (1998). EIN4 and ERS2 are members of the putative ethylene receptor gene family in Arabidopsis. Plant Cell 10: 1321–1332.

17. Ju, C., Yoon, G.M., Shemansky, J. M., Lin, D. Y., Ying, Z. I., Chang, J., Garrett, W. M., Kessenbrock, M., Groth, G., Tucker, M. L., et al. (2012). CTR1 phosphorylates the central regulator EIN2 to control ethylene hormone signaling from the ER membrane to the nucleus in Arabidopsis. Proc. Natl. Acad. Sci. USA 109: 19486–19491.

18. Ju, C., Van de Poel, B., Cooper, E. D., Thierer, J. H., Gibbons, T. R., Delwiche, C. F., Chang, C. (2015). Conservation of ethylene as a plant hormone over 450 million years of evolution. Nat. Plants 1, 14004.

19. Kende, H. (1993). Ethylene biosynthesis. Annu. Rev. Plant Physiol. Plant Mol. Biol. 44: 283–307.

20. Kieber, J. J., Rothenberg, M., Roman, G., Feldmann, K. A., Ecker, J. R. (1993). CTR1, a negative regulator of the ethylene response pathway in Arabidopsis, encodes a member of the raf family of protein kinases. Cell 72: 427–441.

21. Knight, H., Trewavas, A. J., Knight, M. R. (1996). Cold calcium signaling in Arabidopsis involves two cellular pools and a change in calcium signature after acclimation. Plant Cell 8: 489–503.

22. Knight, H., Trewavas, A. J., Knight, M. R. (1997). Calcium signalling in *Arabidopsis thaliana* responding to drought and salinity. Plant J. 12: 1067–1078.

23. Knight, M. R., Campbell, A. K., Smith, S. M., Trewavas, A. J. (1991). Transgenic plant aequorin reports the effects of touch and cold-shock and elicitors on cytoplasmic calcium. Nature 352: 524–526.

24. Knight, M. R., Smith, S. M., Trewavas, A. J. (1992). Wind-induced plant motion immediately increases cytosolic calcium. Proc. Natl. Acad. Sci. USA 89: 4967–4971.

25. Kudla, J., Batistic, O., Hashimoto, K. (2010). Calcium signals: the lead currency of plant information processing. Plant Cell 22: 541–563.

26. Lacey, R. F., and Binder, B. M. (2016). Ethylene Regulates the Physiology of the *Cyanobacterium Synechocystis sp*. PCC 6803 via an Ethylene Receptor. Plant Physiol. 171: 2798–2809.

27. Laothawornkitkul, J., Taylor, J. E., Paul, N. D., Hewitt, C. N. (2009). Biogenic volatile organic compounds in the Earth system. New Phytol. 183: 27–51.

28. Li, W., Ma, M., Feng, Y., Li, H., Wang, Y., Ma, Y., Li, M., An, F., Guo, H. (2015). EIN2-directed translational regulation of ethylene signaling in Arabidopsis. Cell 163: 670–683.

29. Loreto, F., and D’Auria, S. (2022). How do plants sense volatiles sent by other plants? Trends Plant Sci. 27: 29–38.

30. Luan, S., and Wang, C. (2021). Calcium Signaling Mechanisms Across Kingdoms. Annu. Rev. Cell Dev. Biol. 37: 311–340.

31. McAinsh, M. R., Clayton, H., Mansfield, T. A., Hetherington, A. M. (1996). Changes in stomatal behavior and guard cell cytosolic free calcium in response to oxidative stress. Plant Physiol. 111: 1031–1042.

32. McDaniel, B. K., and Binder, B. M. (2012). Ethylene receptor 1 (ETR1) is sufficient and has the predominant role in mediating inhibition of ethylene responses by silver in Arabidopsis thaliana. J. Biol. Chem. 287: 26094–26103.

33. Merchante, C., Brumos, J., Yun, J., Hu, Q., Spencer, K. R., Enríquez, P., Binder, B. M., Heber, S., Stepanova, A. N., Alonso, J. M. (2015). Gene-specific translation regulation mediated by the hormone-signaling molecule EIN2. Cell 163: 684–97.

34. Mount, S. M., and Chang, C. (2002). Evidence for a plastid origin of plant ethylene receptor genes. Plant Physiol. 130: 10–14.

35. Papon, N., and Binder, B. M. (2019). An evolutionary perspective on ethylene sensing in microorganisms. Trends Microbiol. 27: 193–196.

36. Qiao, H., Shen, Z., Huang, S. S., Schmitz, R. J., Urich, M. A., Briggs, S. P., Ecker, J. R. (2012). Processing and subcellular trafficking of ER-tethered EIN2 control response to ethylene gas. Science 338: 390–393.

37. Raz, V., and Fluhr R. (1992). Calcium Requirement for Ethylene-Dependent Responses. Plant Cell 4: 1123–1130.

38. Resnick, J. S., Rivarola, M., Chang, C. (2008). Involvement of RTE1 in conformational changes promoting ETR1 ethylene receptor signaling in Arabidopsis. Plant J. 56: 423–431.

39. Resnick, J. S., Wen, C. K., Shockey, J. A., Chang, C. (2006). REVERSION-TO-ETHYLENE SENSITIVITY1, a conserved gene that regulates ethylene receptor function in Arabidopsis. Proc. Natl. Acad. Sci. USA 103: 7917–7922.

40. Sakai, H., Hua, J., Chen, Q. G., Chang, C., Medrano, L. J., Bleecker, A. B., Meyerowitz, E. M. (1998). ETR2 is an ETR1-like gene involved in ethylene signaling in Arabidopsis. Proc. Natl. Acad. Sci. USA 95: 5812–5817.

41. Sanderson, M. J., Thorne, J. L., Wikström, N., Bremer, K. (2004). Molecular evidence on plant divergence times. Am. J. Bot. 91: 1656–1665.

42. Schaller, G. E., and Bleecker, A. B. (1995). Ethylene-binding sites generated in yeast expressing the Arabidopsis ETR1 gene. Science 270: 1809–1811.

43. Schaller, G. E., Ladd, A. N., Lanahan, M. B., Spanbauer, J. M., Bleecker, A. B. (1995). The ethylene response mediator ETR1 from Arabidopsis forms a disulfide-linked dimer. J. Biol. Chem. 270: 12526–12530.

44. Tian, W., Wang, C., Gao, Q., Li, L., Luan, S. (2020). Calcium spikes, waves and oscillations in plant development and biotic interactions. Nat. Plants 6: 750–759.

45. Wang, W., Esch, J. J., Shiu, S. H., Agula, H., Binder, B. M., Chang, C., Patterson, S. E., Bleecker, A. B. (2006). Identification of important regions for ethylene binding and signaling in the transmembrane domain of the ETR1 ethylene receptor of Arabidopsis. Plant Cell 18: 3429–3442.

46. Ward, J. M., Mäser, P., Schroeder, J. I. (2009). Plant ion channels: gene families, physiology, and functional genomics analyses. Annu. Rev. Physiol. 71: 59–82.

47. Warner, H. L., and Leopold, A. C. (1969). Ethylene evolution from 2-chloroethylphosphonic Acid. Plant Physiol. 44: 156–158.

48. Wen, X., Zhang, C., Ji, Y., Zhao, Q., He, W., An, F., Jiang, L., Guo, H. (2012). Activation of ethylene signaling is mediated by nuclear translocation of the cleaved EIN2 carboxyl terminus. Cell Res. 22: 1613–1616.

49. Yang, S. F., and Hoffman, N. E. (1984). Ethylene biosynthesis and its regulation in higher plants. Ann. Rev. Plant Physiol. 35: 155–189.

50. Zhao, M. G., Tian, Q. Y., Zhang, W. H. (2007). Ethylene activates a plasma membrane Ca(2+)-permeable channel in tobacco suspension cells. New Phytol. 174: 507–515.

